# *THI1* Gene Evolutionary Trends: A Comprehensive Plant-Focused Assessment via Data Mining and Large-Scale Analysis

**DOI:** 10.1101/2023.10.12.562044

**Authors:** Henrique Moura Dias, Naiara Almeida de Toledo, Ravi V. Mural, James C. Schnable, Marie-Anne Van Sluys

## Abstract

Molecular evolution analysis typically involves identifying selection pressure and reconstructing evolutionary trends. This process usually necessitates access to specific data related to a target gene or gene family within a particular group of organisms. While recent advancements in high-throughput sequencing techniques have resulted in the rapid accumulation of extensive genomics and transcriptomics data and the creation of new databases in public repositories, extracting valuable insights from such vast datasets remains a significant challenge for researchers. Here, we elucidated the evolutionary history of *THI1*, a gene responsible for encoding thiamine thiazole synthase. The thiazole ring is a precursor for vitamin B1 and crucial cofactor in primary metabolic pathways. We conducted a comprehensive search for *THI1* information within public repositories with careful curation to achieve this. Our searches reveal an evolutionary trend of 702 *THI1* homologs of Archaea and Eukarya, with a detailed focus on plants. The green lineage of these organisms preserved the THI4 protein domain throughout its diversification by incorporating the N-terminus and targeting chloroplasts. Furthermore, evolutionary pressures and lifestyle appear to be associated with retention of TPP-riboswitch sites and consequent dual post-transcriptional regulation of the *de novo* biosynthesis pathway in basal groups. Multicopy retention of *THI1* is not a typical plant pattern, even successive rounds of genome duplications. Additionally, we identified the diversification of cis-regulatory sites in plants with the conservation of biological processes associated with the initial stages of seed development and preservation of the transcriptional pattern during the diurnal cycle. Our data mining of 484 transcriptome datasets supports this finding and brings a new look at public repositories and evolutionary trends to *THI1*.

## INTRODUCTION

Thiamine is an essential water-soluble vitamin central to various metabolic processes in all living organisms (Yaman *et al*. 2021). All cellular life on Earth utilizes thiamine as a fundamental coenzyme for critical enzymes involved in energy metabolism, including those in the citric acid cycle and the pentose phosphate pathway (Forlani *et al*. 1999; Frank *et al*. 2007; Rapala-Kozik 2011; Rocha *et al*. 2014). The synthesis of thiamine involves two independent pathways culminating in the production of heterocyclic molecules: thiazole and pyrimidine moieties (Kong *et al*. 2008; Jurgenson *et al*. 2009). These will then be condensed into the thiamine moiety to further phosphorylated into TPP (thiamine pyrophosphate). While the core pathway for thiamine biosynthesis is well-conserved across different organisms, including plants, some variations can arise due to evolutionary processes.

In plants, *de novo* thiamine biosynthesis begins with the availability of HMP (hydroxymethyl-pyrimidine) resulting from the conversion of aminoimidazole ribonucleotide (AIR) and S-adenosylmethionine (SAM), catalyzed by phosphomethylpyrimidine synthase (THIC). This pathway is similar to that found in bacteria and archaea (Jurgenson *et al*. 2009). In parallel, NAD+ and glycine are converted to HET (hydroxyethyl-thiazole) by thiamine thiazole synthase (THI1 or THI4) and sulfur from a backbone cysteine in the polypeptide chain itself, a suicide protein. This process is conserved between plants (THI1) and yeasts (THI4) (Chatterjee *et al*. 2009, 2011).

Since thiamine cannot be synthesized *de novo* in animals, including humans, it is necessary to obtain it from external sources such as the diet or environment (Singleton. and Martin 2001). The required amount of thiamine cofactor for growth and development is relatively small on a molar basis (Hanson *et al*. 2016). Moreover, thiamine biosynthesis in plants is energetically expensive due to synthesis and enzymatic turnover costs (Hanson *et al*. 2018). Consequently, the regulatory mechanisms controlling gene expression are complex and involve various factors such as transcription factors, riboswitches, and signaling pathways. These mechanisms allow the organism to fine-tune thiamine biosynthesis based on its metabolic needs and environmental conditions (Ribeiro *et al*. 2005; Wachter *et al*. 2007; Bocobza and Aharoni 2008; Rapala-Kozik *et al*. 2012; Fitzpatrick and Noordally 2021).

This complex regulatory network that governs thiamine biosynthesis involves crucial genes, including the thiamine thiazole synthase gene (*THI1*), responsible for synthesizing the thiazole ring. Understanding the evolutionary significance of *THI1* provides insights into organisms’ adaptive strategies and co-evolutionary dynamics relying on external sources of thiamine and regulatory mechanisms. THI1 is a highly conserved protein containing 346 amino acid (aa) residues with a functional domain THI4 and is present in all plants, fungi, and some archaea (Machado *et al*. 1996, 1997; Godoi *et al*. 2006; Chatterjee *et al*. 2011; Eser *et al*. 2016). In plants, *THI1* genes have been identified and their expression patterns have been studied in model plants like Arabidopsis (Rapala-Kozik *et al*. 2012) and in essential crops such as rice (Wang *et al*. 2016), cassava (Mangel *et al*. 2017), wheat, and barley (Joshi *et al*. 2020), as well as sugarcane (Dias *et al*. 2023).

Molecular evolution and genomics continue to uncover insights into how pathways and genes evolve, providing a better understanding of the origins and variations seen in biochemical processes across different organisms (Jacques *et al*. 2023). We exploited a pipeline incorporating phylogenomic approaches to identify the features of the *THI1* gene in sequenced genomes. Integrated large-scale phylogenetics analysis and transcriptome datasets were used to investigate selection pressures, promoter conservation, expression divergence, and gene underlying duplicate gene evolution. The results of this study lay a substantial foundation for further investigating the contributions of gene evolution and diversification, regulatory process, and adaptative evolution in plants for the *THI1* gene. These results bring information about the essential characteristics of the distribution of THI1 homologs in the Tree of Life, with plants as the main focus of the study, which provides a solid foundation for further investigation of thiamine biofortification through the engineering pathway.

## MATERIALS AND METHODS

### Database Search and THI1 Sequence Mining

To undertake a large-scale search for THI1 protein in the three domains of life, a total of 754 proteins were obtained (Table S1). An investigation by BLASTp and HMMER has been performed with the Arabidopsis THI1 (AT5G54770) protein as the input sequence. Low-scoring hits were filtered, and only sequences with the highest similarity (lowest Evalue and higher bit score) were retained for further analyses. We used Phytozome v13 (https://phytozome-next.jgi.doe.gov/; last accessed January 24, 2023), PLAZA database (https://bioinformatics.psb.ugent.be/plaza/, last accessed December 10, 2021), Ensembl Bacteria (http://bacteria.ensembl.org/, last accessed December 10, 2021), Ensembl Fungi (http://fungi.ensembl.org/, last accessed December 10, 2021) and Ensembl Protists (http://protists.ensembl.org/, last accessed December 10, 2021). They were only maintained as the most representative protein isoform encoded by each unique gene (defined as the most complete isoform). We discard partial sequences and proteins of pseudogenes (often less than 200 residues) by eliminating sequences shorter than 80% coverage of functional domain (THI4) compared to *A. thaliana* THI1 (AtTHI1, 233 aa) and *S. cerevisiae* (SacTHI4, 255 aa).

### Protein Sites Analyses

We used 294 representative species protein sequences from main groups across the Archaeplastida from Streptophyte and Chlorophyte. The amino acid sequences of the THI1 proteins were aligned with MAFFT v7 using the default parameters (Katoh and Standley, 2013). To extract the protein domains, an InterProScan v90 search (Paysan-Lafosse *et al*. 2023) of the InterPro and PfamA protein domain database was performed in the Geneious prime (version 2023.0.1) instance. The domain structures and sequence alignment of the THI1 proteins were visualized with Geneious prime. The N-terminal extension was annotated as described previously (Chabregas *et al*. 2003; Dias *et al*. 2023). To predict the signal peptide cleavage site in N-terminal protein, we used SignalP (Armenteros *et al*. 2019b), and for subcellular localization prediction, we used TargetP (Armenteros *et al*. 2019a). Based on previous studies, the Cys-205 sulfur donor site was annotated in this study (Chatterjee *et al*. 2011; Joshi *et al*. 2020; Dias *et al*. 2023).

### Identification of TPP-riboswitches in *THI1* Sequences

We performed a search for TPP-riboswitches on the entire plant *THI1* CDS (coding sequences) database (Table S4). A typical TPP-riboswitch sequence (RF00059) was used as a search standard by local BLAST against the plant *THI1* CDS database, using an E-value cutoff of 10e-5 and identity ≥ 70%. For further validation of TPP-riboswitches detected by BLAST, we exploited these sequences to see riboswitches using Rfam scanning (Kalvari *et al*. 2021). We also performed the gene search strategy to avoid errors based on the annotation of CDS’s. As we did not find any new TPP-riboswitch, we could confirm the absence of TPP-riboswitch for the sequences that did not have positive results in the first search with CDS.

### Phylogeny Inference

Full-length protein sequences were aligned using MAFFT v7.450 with default parameters (Katoh and Standley 2013). For the construction of the phylogenetic tree, the search for the best model was carried out using the SMS “Smart Model Selection” approach (Lefort *et al*. 2017) in the “ATGC Montpellier Bioinformatics Platform” (http://www.atgc-montpellier.fr/) with the aid of the PhyML v3.0 tool (Guindon *et al*. 2010), following the analyzes with the JTT+R model of amino acid substitution (see Supplementary file 1). The tree topology was modeled by the NJ (Neighbor-Joining) method, where the statistical support of the arms was calculated by the aLRT-SH method (Lefort *et al*. 2017).

### Identification of *THI1* Synteny Blocks

The *THI1* synteny block was identified using genome collinearity in the PLAZA database tool (https://bioinformatics.psb.ugent.be/plaza/; Van Bel *et al*. (2022)). For checking, we performed the chromosome-level sequence alignments using CoGe (https://genomevolution.org/coge/SynMap.pl; Lyons and Freeling (2008); Lyons *et al*. (2008)) in pairwise comparisons (see Table S3 for details).

### Cis-regulatory Element Prediction and Characterization

We performed a comparative analysis of the promoter regions of the different occurrences of *THI1* in representants of Archaeplastida, specifically Embryophytes (Eudicots, Monocots, Gymnosperms, Lycophytes, and Bryophytes), and core Chlorophyte following a methodology proposed in a previous study (Dias *et al*. 2023). Upstream sequences of 1,500 bp from the start codon were assessed for conserved features and motifs using MEME suite tools (https://meme-suite.org/meme/; Bailey *et al*. (2009)). We used all other default MEME parameters except for 20 motifs from 5 to 25 bp in length, with an E-value 0.05 (Powell *et al*. 2019). The retrieved motifs were run through TomTom (Gupta *et al*. 2007) via the JASPAR Core Plants 2023 database (Khan *et al*. 2018), and the respective UniProt IDs results were collected if the p-value 0.01. The UniProt IDs were used to collect biological GO terms for the function assignment of each motif. g:Profiler (Raudvere *et al*. 2019) was used to analyze the over-representation of GO terms. An R package ggplot2 (Wickham 2016) was used to generate a heatmap with adjustments in Inkscape Illustrator.

### Transcriptome Data Compilation and Analysis

RNAseq data of different samples from 11 species (*C. reinhardttii, P. patens, S. moellendorffii, O. sativa, B. distachyon S. bicolor, Z. mays, S. italica, A. thaliana, G. max*, and *S. lycopersicum*) were grouped into three different classes: organs/tissues, rhythmics cycle, and seed development. A total of 484 different RNA-seq samples were used. These datasets are publicly available and were downloaded from Expression Atlas (https://www.ebi.ac.uk/gxa/experiments) and ePlant BAR (https://bar.utoronto.ca/#GeneExpressionAndProteinTools). For more details, see Table S7.

## RESULTS AND DISCUSSION

Despite its recognized importance, numerous aspects surrounding this gene remain enigmatic, evoking scientific questions regarding the loss of the pathway in animals and its maintenance in plants and fungi. Also, the thiazole protein domain is highly conserved despite its patchy distribution in cellular organisms. Here, an integrative approach focused on mapping the presence of *THI1* in well-resolved phylogenetic trees in fully sequenced genomes is presented. The distribution and classification of all THI1 homologs based on protein structure, presence/absence of TPP-riboswitches, promoter gene structure, and inference of gene regulatory mechanisms based on expression patterns constituted a compilation of information in available databases shown in this comprehensive study.

### Identification and Distribution of THI1 in the Three Domains of Cellular Life

With the increasing availability of fully sequenced genomes, a comparative genomic approach can now be employed to analyze genes across a broad range of species. To investigate THI1, we searched public databases for sequences from the three domains of life (Eukarya, Archaea, and Bacteria), using *THI1* genes previously cloned from *A. thaliana* and *S. cerevisiae* (Machado *et al*. 1996, 1997) as references. Based on a BLASTp and HMMER search against protein databases of the three domains of life, all gene entries from fully sequenced and annotated genomes are presented (Table S1).

Our scientific investigation revealed exciting insights into the distribution of THI1 sequences among different organisms. Among Eukaryotes, Fungi showed the highest abundance of THI1, with 428 sequences found across 421 species. Following closely were Viridiplantae, with 216 sequences present in 108 species. Chlorophyta had ten sequences distributed among six species, while Oomycota had 13 species encompassing three orders: Saprolegniales (eight members), Pythiales (five members), and Peronosporales (two members). Two species in Alveolata presented THI1 sequences. Additionally, a single sequence was identified in Cryptophyceae, which belongs to non-green algae.

In the Archaea domain, most THI1-harboring species belong to the Euryarchaeota and Thaumarchaeota, totaling 30 sequences distributed across 18 species. Remarkably, THI1 was absent in Bacteria, providing evidence for a distinct biochemical pathway for HET synthesis from Archaea and Eukarya (Figure 1A). Consequently, our findings strongly support the hypothesis that THI1 fulfills a unique functional role in Archaea and Eukarya. However, the question of its origin remains intriguing. In this context, the analogous pathway observed in bacteria for the thiazole ring formation is noteworthy. In bacteria, HET biosynthesis follows a four-step path involving several enzymes, namely ThiF, ThiS, ThiG, and ThiO, as reported (Begley *et al*. 1999; Hazra *et al*. 2009).

**Figure 1.**
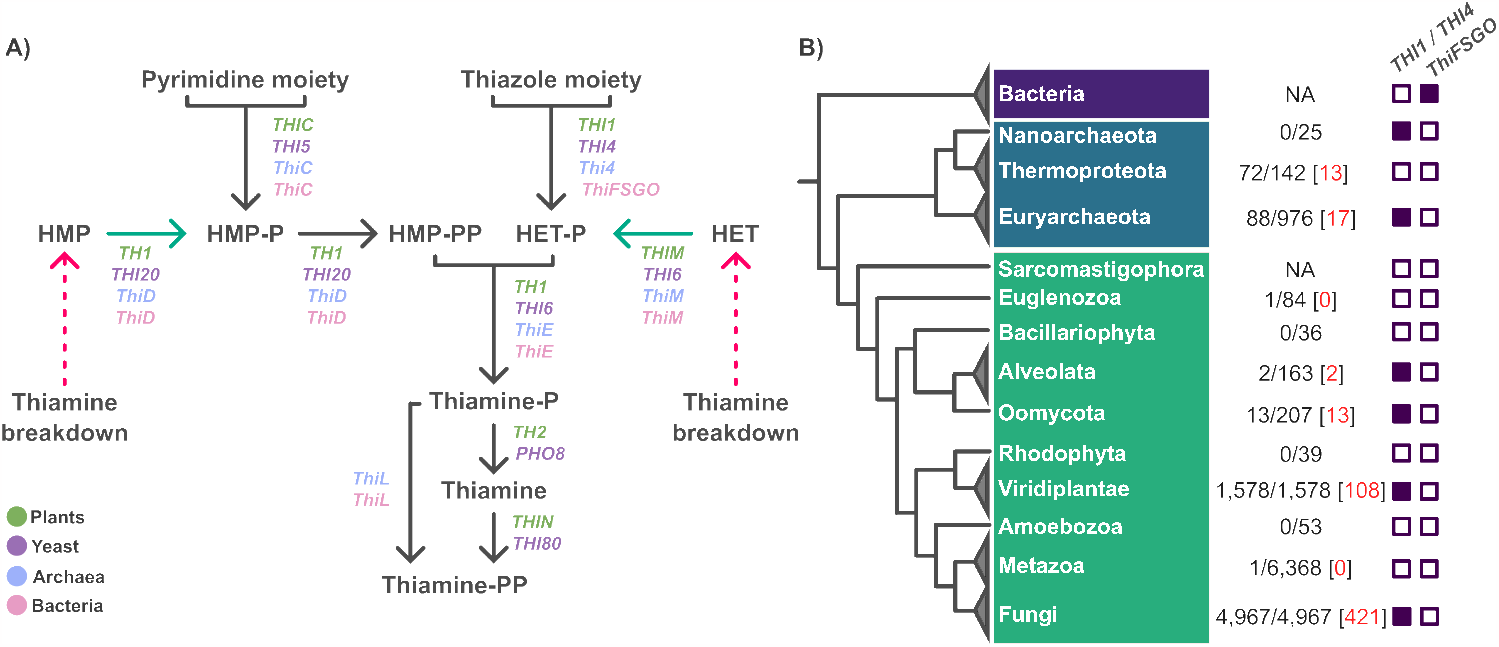
Identification of *THI1* homologs along Tree of Life. **A)** The thiamine KEGG pathway is summarized for Archaea, Bacteria, and Eukarya (represented by yeast and plants). In Archaea, genes involved in thiamine biosynthesis (*Methanocaldococcus jannaschii*) are shown in blue, whereas homologs in Bacteria (*Bacillus subtilis*) are indicated in pink. Eukarya is represented by purple (yeast, *Saccharomyces cerevisiae*) and green (plants, *Arabidopsis thaliana*). The dashed line represents the breakdown thiamine pathway, the green line represents the salvage pathway, and the black line indicates the *de novo* biosynthesis pathway. **B)** Scattered and restricted distribution of *THI1/THI4* homologs on a representative Tree of Life phylogenetic tree. The number of *THI1/THI4* homologs was mapped onto a representative phylogenetic tree consisting of major species groups (according to NCBI Taxonomy). At the level of each group, the ratio of the number of homologs to the total number of species deposited in the NCBI. The red numbers represent the *THI1* homologs found on the database used in the study (see Table S1 and Material and Methods).

Additionally, it is important to note that the determination of the presence of *THI1* genes in specific taxa was limited by their genomes’ incomplete or draft nature. Despite this challenge, the widespread and diverse distribution of *THI1* across different domains of life inspired us to map the identified *THI1* genes on a phylogenetic tree consisting solely of species with complete genome sequences. This mapping allowed us to present, at each taxonomic level, the ratio between the number of *THI1* genes and the total number of species included in our study (refer to Figure 1B and Table S1 for detailed information). To date, we have not identified evidence of *THI1* in complete animal genomes, but we have identified one report of evidence of horizontal transfer in nematodes (Craig *et al*. 2009). While constrained by the unavailability of the Heterodera glycines genome, this study underscores the hypothesis that the THI1 gene has been lost in animals, with instances of horizontal transfer occurring only in rare and exceptional cases. By employing a phylogenomic approach, our study sheds light on the evolutionary history of THI1, revealing significant diversification and specialization within this protein. These findings provide valuable insights into how THI1 has evolved and adapted to different organisms throughout evolution, which will be presented in the following sections.

### Phylogenetic Analysis and Diversification of *THI1* Homologs Across Eukarya and Archaea Show the Emergence of Thiamine Thiazole Synthase Targeting

In our study, a phylogenetic analysis of the THI1 containing only PF01946 (THI4; Q38814) results in an unresolved tree topology that still needs to be reached, as depicted in Figure S1. Therefore, a second round of analysis was performed using the entire THI1 protein sequence, which support its presence in Archaea and Eukarya, with an uneven distribution among closely related lineages (Figure 2A). The phylogenetic tree topology revealed that Archaea branches out from Eukarya, where THI1 is only found in five major groups: Fungi, Oomycota, Embryophytes, Chlorophyte, and two Alveolates (Dinophyceae and Perkinsoza). Fungi sequences are more diversified compared to the remaining lineages. Chrlorophyta and Streptophyta are clearly non-monophyletic and the two clusters close to Oomycota and Alveolates.

**Figure 2.**
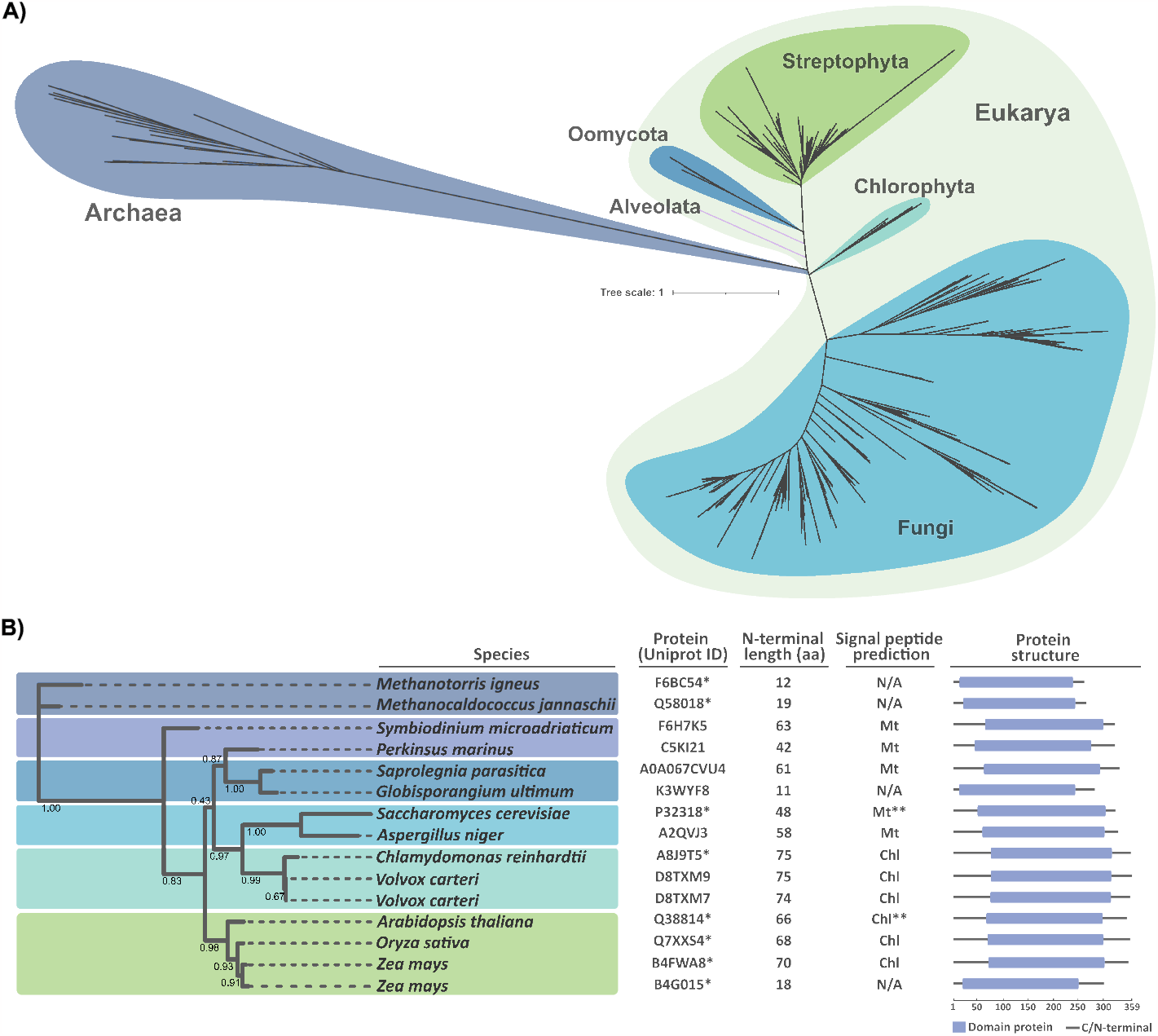
Phylogeny and features of THI1 proteins in the three domains of life. **A)**. Phylogenetic unrooted trees were constructed based on the amino acid sequences of 853 selected THI1 proteins using the maximum likelihood method with 1,000 bootstrap replicates. **B)** Representative THI1 protein homologs characterized in the literature are shown, including the experimentally characterized proteins marked with an asterisk (Eser *et al*. 2016; Singleton 1997; Croft *et al*. 2007; Chabregas *et al*. 2001; Woodward *et al*. 2010; Wang *et al*. 2006), and subcellular localization confirmed is marked with two asterisks (Chabregas *et al*. 2003; Chatterjee *et al*. 2011). Domains were visualized with Geneious Prime v.3.22. Signal peptide prediction is classified into three groups: Chloroplast transit peptide (Chl); Mitochondrial transit peptide (Mt); N/A, not applicable. The scale bar beneath the domain illustration shows each protein’s amino acid (aa) length.

Our results support the potential use of the N-terminal region of the gene as a DNA barcode. The unrooted tree displayed a significant divergence between Archaea and Eukarya which is supported by the molecular clock estimates and phylogenetic analyses indicating that the separation of Archaea and Eukaryotes from their last common ancestor (LCA) roughly by 2 billion years ago (Cox *et al*. 2008; Eme *et al*. 2017).

We further investigated the protein structure and annotated the proteins for their THI4 domain extension, size, and amino acid composition for the best-annotated sequence from Archaea and Eukarya subclades (Figure 2B). Our study shows that the domain is conserved in all amino acid sequences sampled, with the size of the N-terminal portion varying from 12 to 70 aa. The N-terminus is an added novelty to each THI4 domain. Furthermore, the presence of a transit peptide is only present in Eukarya, mainly predicted as mitochondrial targeting in Alveolates, Oomycota, and Fungi, whereas in plants, targeting to chloroplasts is a consensus (Dias *et al*. 2023). The Archaea versions do not possess extended N-terminal sequences, as no organelle targeting is needed (Figure 2B).

The compartmentalization and development of the eukaryotic endomembrane system (Bohnsack and Schleiff 2010) enforced the adaptation of the ancestral protein to acquire a translocation system, as seen in the case of THI1. Furthermore, most eukaryotic translocation systems can be traced back to those developed very early in the evolution of life (Patron and Waller 2007). Hence, the evolutionary analysis of the THI1 protein encompassing the N-terminal peptide not only identifies the diversification of large groups in the Tree of Life but also provides information about the compartmentalization of the thiamine biosynthesis compared to cells from Archaea and Bacteria. The investigation into the evolutionary timeline of the thiazole pathway has unveiled unexpected revelations about the development of compartmentalized eukaryotic metabolism (Lunn 2007). Notably, the protein THI1 is directed to mitochondria and chloroplasts in plants through a post-transcriptional mechanism, as demonstrated by (Chabregas *et al*. 2001). A question remains: What selective constraints drove compartmentalization, and which pathways?

### Ancestry of the *THI1* Gene in Green Plant Lineages

To investigate the extent of THI1 diversity among plants, we conducted a thorough search for *THI1* homologs in the complete genomes of 148 plant species. Subsequently, we constructed an unrooted maximum likelihood phylogenetic tree based on the global alignment of amino acid sequences (Figure 3). To further enhance our understanding of thiamine thiazole synthase evolution in Viridiplantae, we annotated each gene’s genomic sequence length, CDS and protein size, intron number, and transcript splicing sites, as summarized in Table S1.

**Figure 3.**
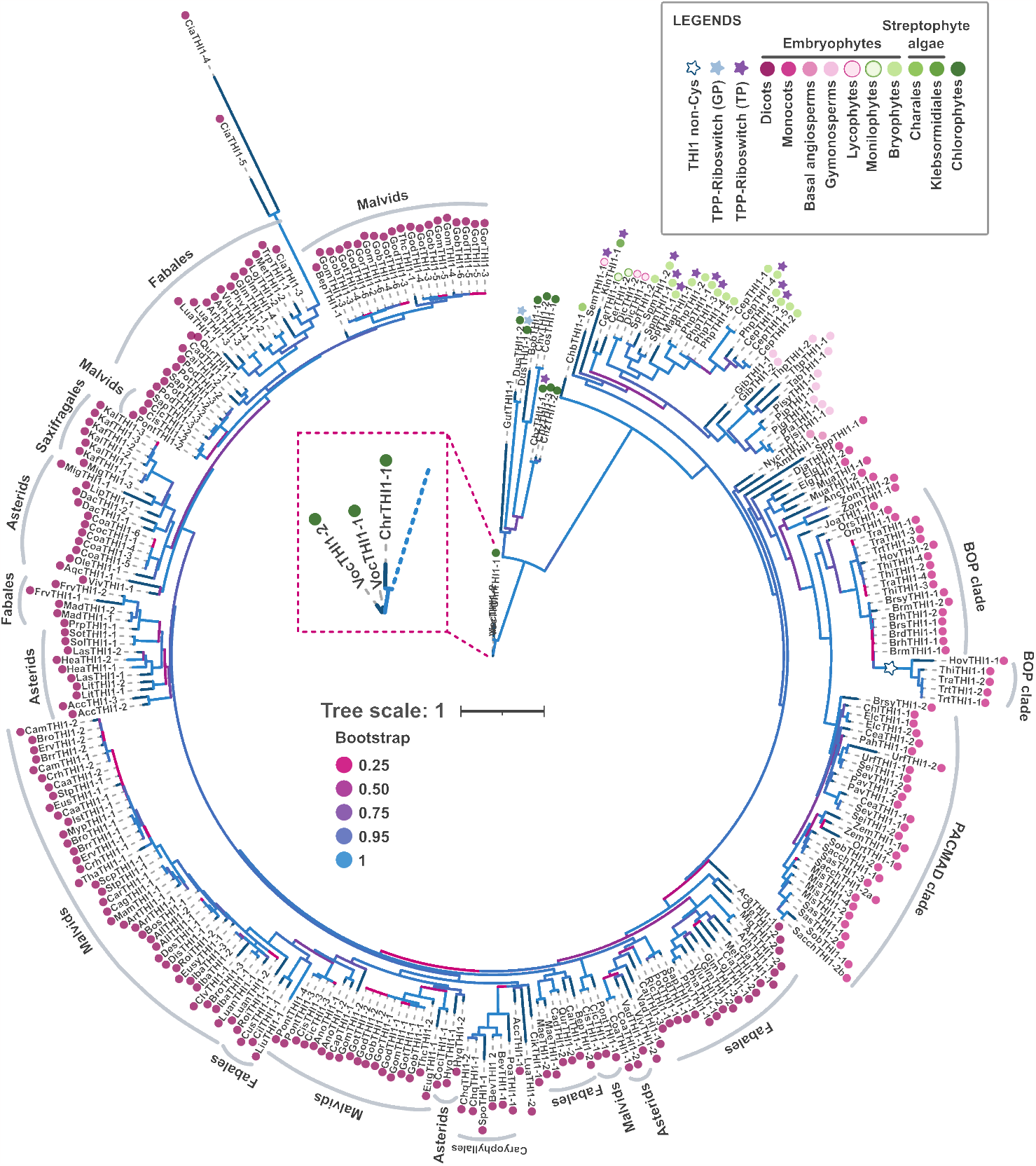
Phylogenetic tree of THI1 in plants. The phylogenetic tree was constructed based on the amino acid sequences of 294 THI1 proteins using the maximum likelihood method with 1,000 bootstrap replicates. Numbers at nodes are branch support values estimated by the aLRT SHlike method implemented in PhyML3.0 (see Methods). Colors on the branches indicate bootstrap values. The presence of a riboswitch is indicated with a blue star (genomic detection) or pink star (transcript detection). A blue star represents the diversity of Cysteine in the catalytic site of the sulfur donor (see Joshi *et al*. (2020); Dias *et al*. (2023)).

To gain insight into the evolutionary history of THI1 in plants, we used Chlorophyta as an outgroup to the Streptophyta lineage. These findings support the arrangement of these proteins into 3 primary clades. The first clade, representing the ancestral lineage, comprises core chlorophytes like Chlorophyceae and Trebouxiophyceae. The second clade encompasses streptophyte algae such as Charophyceae and Klebsormidiales. The third clade includes a diverse array of embryophytes—Bryophytes, Monilophytes, Lycophytes, Gymnosperms, Basal angiosperms, Monocots, and Eudicots—thus often referred to as Land Plants (Figure 3).

Undoubtedly, the limited diversity of available genomes has hindered our ability to make robust evolutionary inferences for underrepresented groups of non-seed plants, as noted (Rensing 2017). Nevertheless, the phylogenetic tree has confirmed the expected grouping of THI1 homologs in Bryophyta, Monilophytes, and Lycophytes as sister clades (Puttick *et al*. 2018). The results of this study suggest that THI1 in plants can be traced back to a single ancestral gene, which likely originated from a common ancestor shared between core streptophytes, embryophytes. While Chlorophyta (core Chlorophytes) forms a monophyletic outgroup of Streptophyta (Lewis and McCourt 2004), our analysis, which was focused on thoroughly curated genomes, did not identify *THI1* homologs in other algae members beyond Chlorophyceae (*Chlamydomonas reinhardtii, Chromochloris zofingiensis, Dunaliella salina*, and *Volvox carteri*) and Trebouxionophyceae (*Botryococcus braunii* and *Coccomyxa subellipsoidea*). The restricted distribution of *THI1* genes in certain green algae of core chlorophytes prompted us to search for THI1 homologs in publicly available scaffold assembly in the database PhycoCosm (Grigoriev *et al*. 2021).

Despite the scarcity of assembled and curated green algal genomes, a tBLASTn search employing the THI1 (ChrTHI1) protein of *C. reinhardtii* as a query and setting an E-value threshold of ≤10e-30 and a minimum identity of ≥65% unveiled a consensual occurrence of THI1 homologs in all representatives of core chlorophytes, streptophytes, and land plants. Although these scaffolds cannot facilitate detailed analyses, our search allowed the identification of the distribution of new THI1 homologs in 41 species of green algae (chlorophytes) and ten species in streptophyte algae, by the green algae phylogenetic relationship suggested (Leliaert *et al*. 2012) (refer to Table S1-2).

The comprehensive search for THI1 homologs in green algal lineages accentuates the deep correlation of this gene with plant evolution and diversification. It is widely accepted, as noted (Wodniok *et al*. 2011), that land plants (embryophytes) evolved from streptophyte algae. Our results establish that the origin of THI1 in green lineages predates the diversification of the group of embryophytes, streptophytes, and chlorophytes, potentially one billion years ago in the Neoproterozoic (Parfrey *et al*. 2011), which suggests that the formation of the thiazole ring is ancient and conserved by the pathway of thiamine thiazole synthase (THI1) in chloroplasts compartment in green lineages (Figure 2B).

### Post-Transcriptional Dual Control by TPP-riboswitches is Lost in Seed Plants

The relationship between *THI1* homologs from non-seed plants and green algae is further reinforced when considering the retention of TPP-riboswitch motifs in transcripts. Our analysis of the *THI1* homologs revealed that Chlorophytes contain a TPP riboswitch in their respective 5-′UTR based on sequence homology, while Embryophytes contain a TPP-riboswitch in their respective 3-′UTR (Figure 4A). Despite the divergent position of the TPP-riboswitch sites, a detailed inspection of secondary structure content showed that the nucleotides display high conservation of junctions and helices (Figure 4B-C), conserving the structure, although differences were found in length and position.

**Figure 4.**
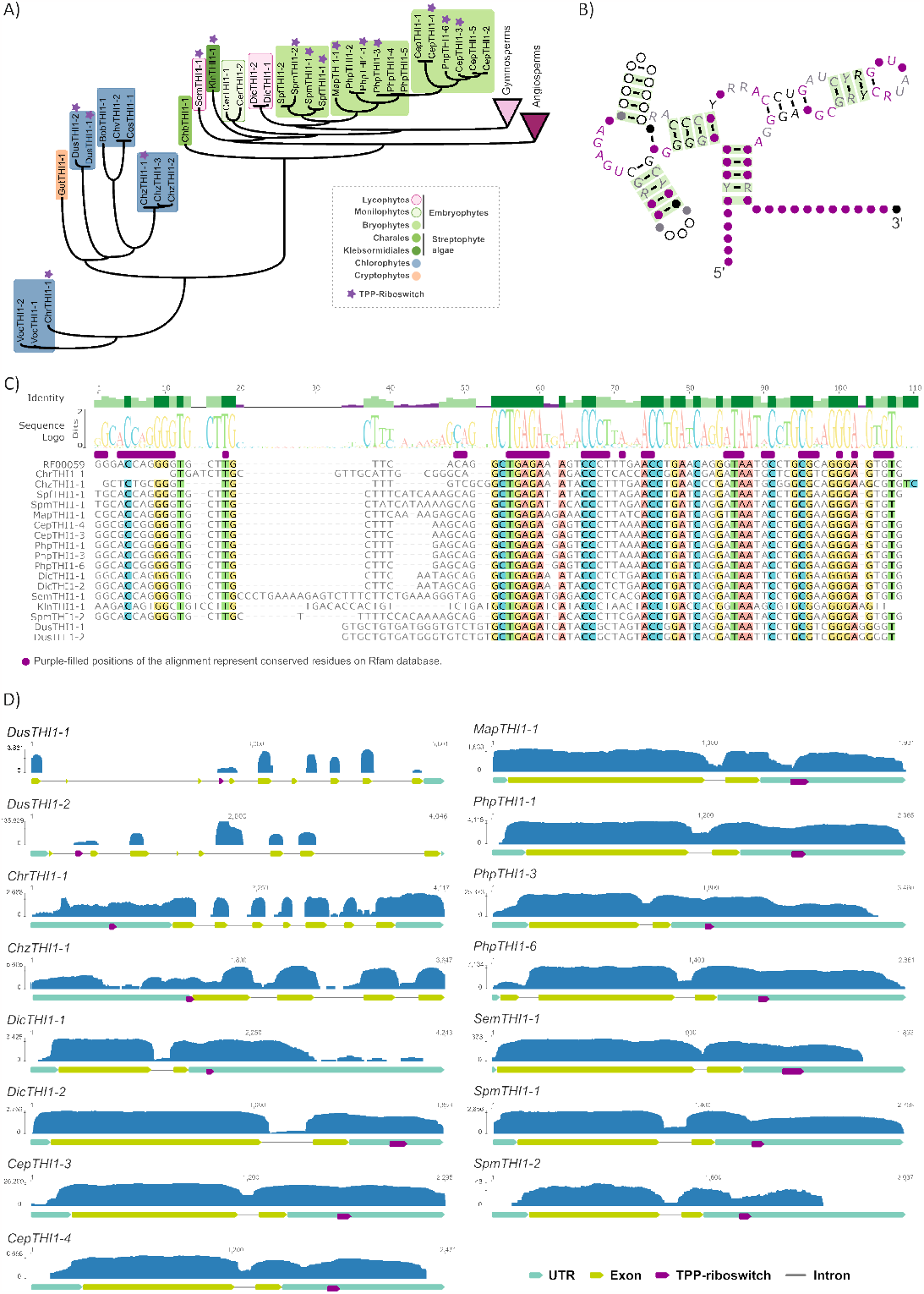
Identification, conserved sites and secondary structure predictions of the TPP-riboswitch (THI element or THI-box). **A**. Phylogeny summary and species harboring THI element in *THI1* homologs. **B)** Consensus secondary structure of TPP-riboswitch aptamer based on Rfam repository (RF00059). Green highlights represent sequence statistically significant base pair with covariation; purple dots are 97% conserved nucleotide; black dots are 90% conserved nucleotide; grey dots are 75% conserved nucleotide; and the with dots are 50% conserved nucleotide. **C)** Multiple alignments of THI-box sequences predicted in 17 *THI1* homologs with annotation of conserved nucleotides (purple boxes). **D)** The read mapping was conducted using a Genious tool with default parameters. The BAM file generated by read mapping was used for visualization. Check section Data Availability to access BioProject ID of RNAseq datasets.

Furthermore, we confirmed that the TPP-riboswitch motif found in our sequence samples is not just a genomic artifact but is transcribed (see Figure 4D). As documented, the thiamine biosynthesis pathway is omnipresent in most plant species (Goyer 2010; Dias *et al*. 2023). However, only a handful of biosynthesis genes are subject to regulation by TPP-riboswitch motifs in various plant species (Figure 1A). Although TPP-riboswitch-based gene regulation is widespread in the *de novo* biosynthesis genes such as the THIC (Bocobza *et al*. 2007; Wachter *et al*. 2007; Moulin *et al*. 2013; McRose *et al*. 2014), it has not been retained in seed plants, only a few representatives of non-seed plants harbor the post-transcriptional regulation in *THI1* (Bocobza *et al*. 2007; McRose *et al*. 2014) (Figure 4A).

It is worth noting that a cluster (Bootstrap >0.90) consisting of THI1 homologs Gymnosperm, Bryophytes, Monilophytes, Lycophytes, and basal angiosperms (*Nymphaea colorata* and *Amborella trichopoda*) (Figures 3 and 4A) represents the Angiosperms where TPP-riboswitch motifs are absent from all studied *THI1* homologs. These findings support that TPP-riboswitch in *THI1* was lost in the ancestor of seed plants. This dual regulation of the thiamine pathway by TPP-riboswitches in the *THI1* and *THIC* transcripts suggests an essential regulation in green lineages that colonized environments with high humidity, but that is subsequently lost in one branch of the pathway in the thiamine thiazole synthesis moiety and maintained in the pyrimidine branch.

### Only a Few Lineages Retain Multiple *THI1* Paralogs After WGD Events

We observed apparent duplication events at the ancestral nodes of mosses and eudicots (Figure 3 and more resolution in Figure 5). Interestingly, we also observed large-scale duplication events at the species level in eudicots, resulting in diversified *THI1* genes in corresponding species. Notably, *THI1*-harboring monocot members formed two distinct monophyletic subclades, corresponding to the BOP and PACMAD sister groups. In line with another study (Dias *et al*. 2023), *THI1* monocot paralogs formed a well-supported clade, indicating that these genes duplicated more recently than the divergence of monocots and eudicots (Figures 4-5).

**Figure 5.**
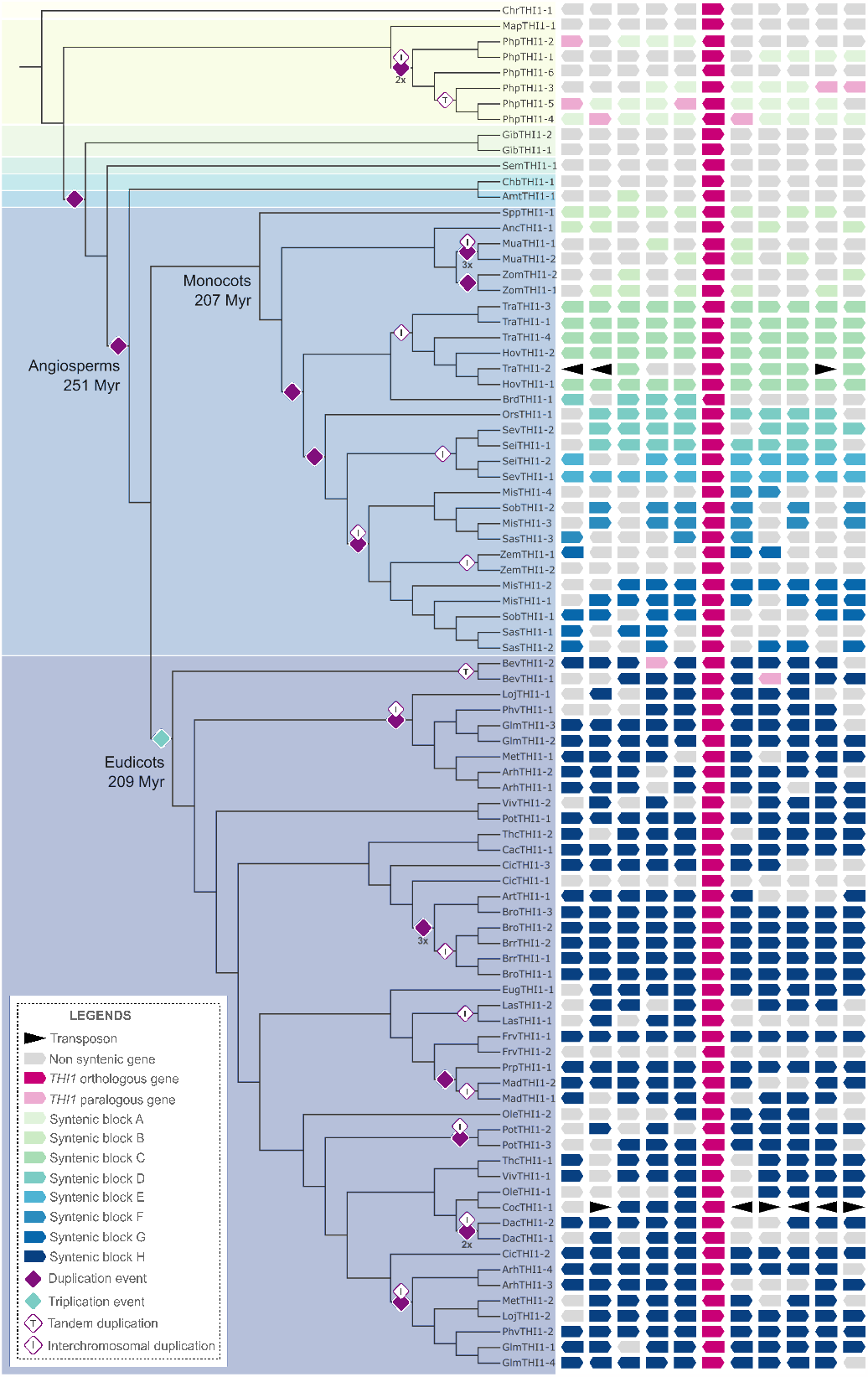
The genomic environment of *THI1* along plant lineage. Phylogenetic representation of plant species sampled in the analysis, distribution of the number of gene duplications, and annotation in millions of years (Ma) belonging to the separation of the main groups of plants (Schneider *et al*. 2004). The synteny of *THI1* is shown following the phylogeny simplified at the chromosomal-level genome assemblies. The PLAZA synteny analysis was manually curated (see Table S3) with adjustments in Inkscape Illustrator. The abbreviations for the plant lineages can be found in Table S3.

In contrast to monocots, eudicots exhibit a divergent *THI1* subclade that comprises numerous subgroups composed of *THI1* paralogs belonging to the Asterids, Caryophyllales, Fabids, Malvids, Vitales, Saxifragalles, Laureales, and Ranunculales. The THI1 paralogs in eudicots are not clustered according to species or family specificity, as observed clearly in Fabids (e.g., *Glycine max, Lotus japonicus*, and *Medicago truncatula*) and Malvids (e.g., *Gossypium spp*., *Carica papaya*, and *Theobroma cacao*). These findings suggest that the duplication or expansion of *THI1* genes in eudicots occurred in ancestors of specific families (see dating of duplicates in Figure 5; (Schneider *et al*. 2004)). However, the retention of multiple paralogs constitutes a recent hybridization event between two closely related species, leading to polyploidy in some crop species, such as *Gossypium sp*. and *Glycine max*. An important point is the variations between paralog sequences and synteny conservation, which support this observation (Figure 5). *THI1* genes showed a complex evolutionary history with several lineage-specific duplication and loss events based on these phylogenetic inferences.

As highlighted by previous studies (Panchy *et al*. 2016; Qiao *et al*. 2019), plant genomes exhibit high rates of evolution, resulting in greater genome diversity and an increased number of gene families derived from duplication events. Although most duplicates are lost following whole-genome duplication (WGD), those gene families that are deemed essential for plant fitness are more likely to be retained, as stated (Panchy *et al*. 2016; Wu *et al*. 2020). To understand the possible duplication mechanism under *THI1*, we explored the retention of the duplicated copies. First, a synteny-based approach was used after retrieving syntenic gene pairs (between and within genomes) using genome collinearity in the PLAZA database tool (Van Bel *et al*. 2022). The accession numbers of the neighborhood genes are listed in Table S3.

Our analyses suggest a scenario where few genomic regions containing the *THI1* gene are maintained and diversified in land plants. These gene pairs and syntenic blocks are represented in Figure 5, in which the representation was manually optimized to depict the eight major *THI1* syntenic blocks and highlight their plant groups. We retrieved a total of 83 *THI1* homologs in plants, with 18 species harboring a single *THI1* homolog, 34 species harboring two *THI1* homologs, and some species showing up to six *THI1* homologs, such as the case of *Physcomitrium patens* (6), *Glycine max* (4), *Arachis hypogaea* (4), *Brassica oleraceae* (3), *Miscanthus sinensis* (4), *Saccharum sp*. (4), *Triticum sp*. (4) (see Table S1 for details).

Accordingly, we assumed that the retention of at least one copy of the *THI1* genes in plants after events of WGD and fractionation (gene loss) (Flagel and Wendel 2009) suggests a critical adaptative feature in the survival and maintenance of plant homeostasis (Papini-Terzi *et al*. 2003; Woodward *et al*. 2010). The expansion of the *THI1* gene is impacted by both whole genome duplication (WGD) and tandem duplication (TD), and the evolutionary history of each plant genome plays a significant role in shaping the *THI1* landscape. Results indicate that the synteny of *THI1* homologs is not conserved across all plant genomes except for eight syntenic blocks that show remarkable conservation (blocks A-H, Figure 5). Interestingly, the genomic neighborhood of plants containing a single *THI1* locus did not exhibit synteny conservation among different groups of embryophytes.

Instead, the genes surrounding the single *THI1* locus displayed substantial variation with a unique combination of genes between distinct plant groups (Figure 5). This supports the hypothesis that *THI1* has an ancestral origin in the plant lineage, with different gene rearrangements around the *THI1* locus. One possible explanation is that duplications generating other *THI1* loci in various plant lineages occurred after the separation of some groups, as observed in Figure 5 (dating made by (Schneider *et al*. 2004). However, evidence suggests that many plant groups share at least one round of WGD (Clark and Donoghue 2018). In this context, the conserved synteny between a single genomic region in multiple blocks may be explained by one or more shared duplications followed by lineage-specific re-diploidization or diploidization events, as proposed (Wendel 2015; Qiao *et al*. 2019).

### *THI1* Genes Reveal a Cluster of Cis-regulatory that Suggests a Conserved Expression Profile Among Plants

In this section, promoter rearrangement was more broadly explored. We first identified 19 motifs along 1,500 bp upstream to ATG of 52 homologs (including paralogs), representing 25 plant species (Table S5; see section methods). Subsequently, we investigated the distribution and retention of these motifs. We identified the top two motifs (Motif-1 and Motif-6) present in all 52 sequences with different degrees of conservation, showing a statistically significant association (E-value ≥ 0.05). Among the 19 motifs, a few overlaps were observed, indicating that throughout the evolution of plants, the *THI1* genes were subject to high rates of mutations in the promoter region compared to the nucleotide coding to the THI4 domain (Figure 6A and Figure S1), revealing the diversity of motifs among plant species.

**Figure 6.**
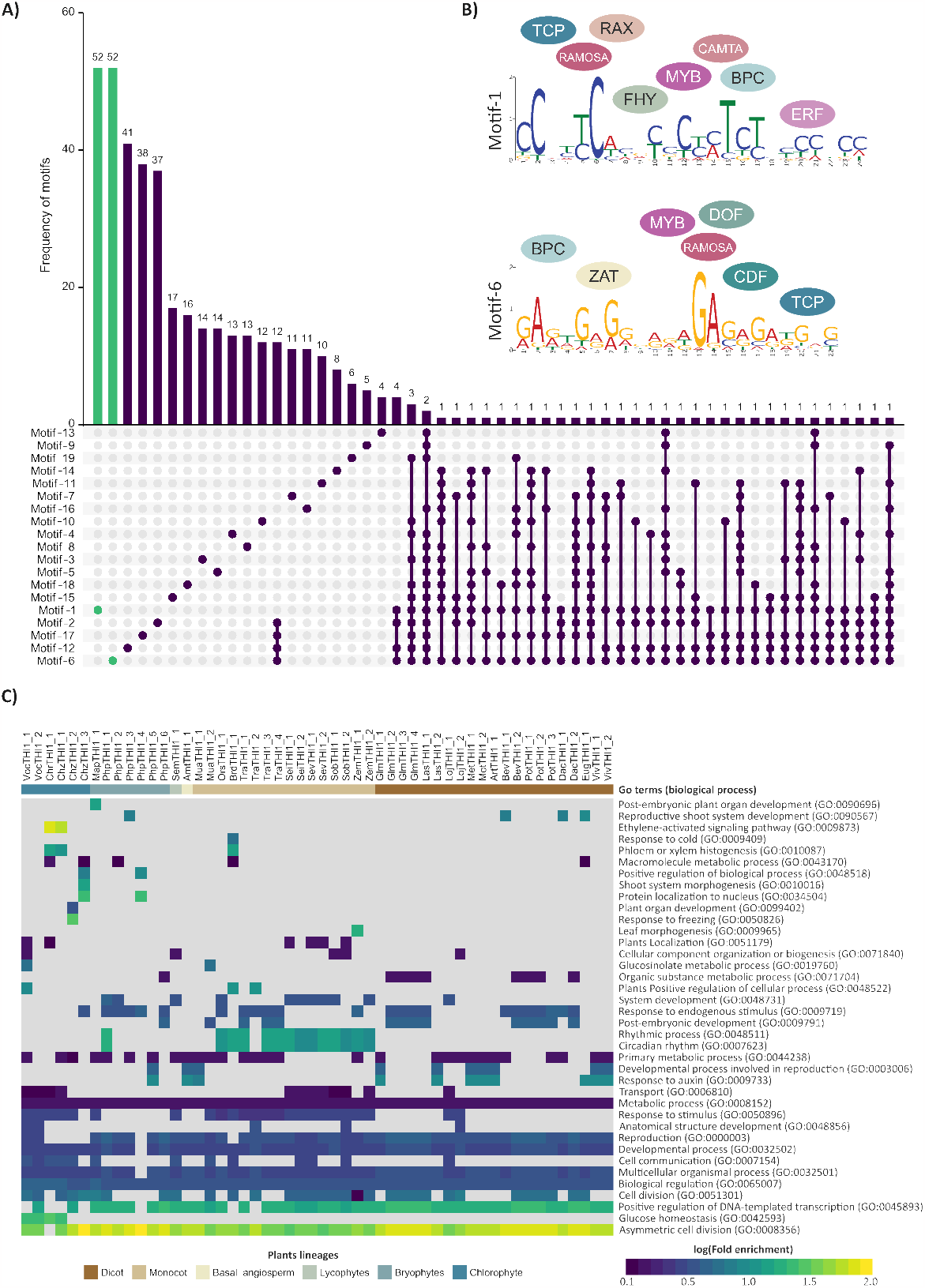
Promoter analysis and motif enrichment of *THI1* homologs. **A)** Compilation and overlap of Motifs in promoter regions (1,500 bp upstream of ATG) at *THI1* homologs. **B)** Transcription Factors (TFs) identified with high association to cis-regulatory elements for Motif-1 and Motif-6. **C)** GO enrichment analysis of motifs localized in the promoter region of *THI1* of representative plants (including core chlorophyte, bryophytes, lycophytes, basal angiosperm, monocots, and eudicots).

Interestingly, both conserved motifs show over-retained cis-regulatory elements sites capable of binding to transcription factors (TFs) such as TCP, RAMOSA, MYB, and BCP (Gallavotti *et al*. 2010; Wang *et al*. 2011; Manassero *et al*. 2013; Li *et al*. 2022), mainly associated with regulation of developmental stages. In addition, the specific TFs related to Motif-1 (RAX, FHY, CAMTA, and ERF) (Muller *et al*. 2006; Feng *et al*. 2020; Xiao *et al*. 2021) and to Motif-6 (ZAT, DOF, and CDF), also show involvement in the diverse developmental process and stresses adaptation (Figure 6B and Table S5). We assumed that the low conservation of promoter region (Figure S1) and, consequently, reduction of overlap among motifs could reveal functional overlap of TFs, since different TFs present specific binding sites, however, they can respond to a set of phenomena of the same nature (see the reviews of (Bemer *et al*. 2017; Lai *et al*. 2020)). In this case, we used *in silico* approaches to analyze the potential biological process associated with TFs capable of binding the site diversity on the promoter region (see section methods).

Our Gene Ontology (GO) analysis revealed significant enrichment and conservation of GO terms associated with asymmetric cell division (GO:0008356) and positive regulation of DNA-templated transcription (GO:0045893), both showing a log-fold enrichment of >1.2. Additionally, we observed a lack of enrichment but conservation for GO terms related to biological regulation (GO:0065007), multicellular organismal processes (GO:0032501), and developmental processes (GO:0032501) (Figure 6C). The interplay among transcription factor families not only expands the molecular toolkit within plants but also complements TPP-riboswitches, providing supplementary mechanisms to finely regulate the dynamic expression patterns of the *THI1* gene in response to changing environmental conditions. This orchestration ensures precise control of gene expression during crucial stages of plant growth and development, a process previously identified in other vitamin pathways (Strobbe *et al*. 2018). This coordination is essential to maintain the integrity of the pathway. In addition, its understanding it opens up new avenues for engineering synthetic promoters (Jores *et al*. 2021).

### Global view of *THI1* Gene Expression from 484 Transcriptomes

A transcriptomic approach was taken to establish where and when the *THI1* homologs were expressed. We constructed gene expression atlases for nine phylogenetically representative species, including sampling tissues such as aerial part (comprising whole leaves), stem (comprising the sample without leaves or lateral meristem), apical meristem (when present), reproductive organ (comprising flowers of floral tissues or analog tissue) and roots (or analog tissue) (Supplementary Table 7). These include the bryophytes *Physcomitrium patens*, the lycophyte *Selaginella moellendorffii*, the monocots *Oryza sativa, Brachypodium distachyon, Sorghum bicolor* and *Zea mays*, and the eudicots *Arabidopsis thaliana, Solanum lycopersicum* and *Glycine max*. The atlases were constructed by combining publicly available RNA sequencing (RNAseq) data with 484 files generated by independent groups, curated and joined in eFPlant BAR (Waese *et al*. 2017), Expression Atlas (Papatheodorou *et al*. 2018) (Figure 7A).

**Figure 7.**
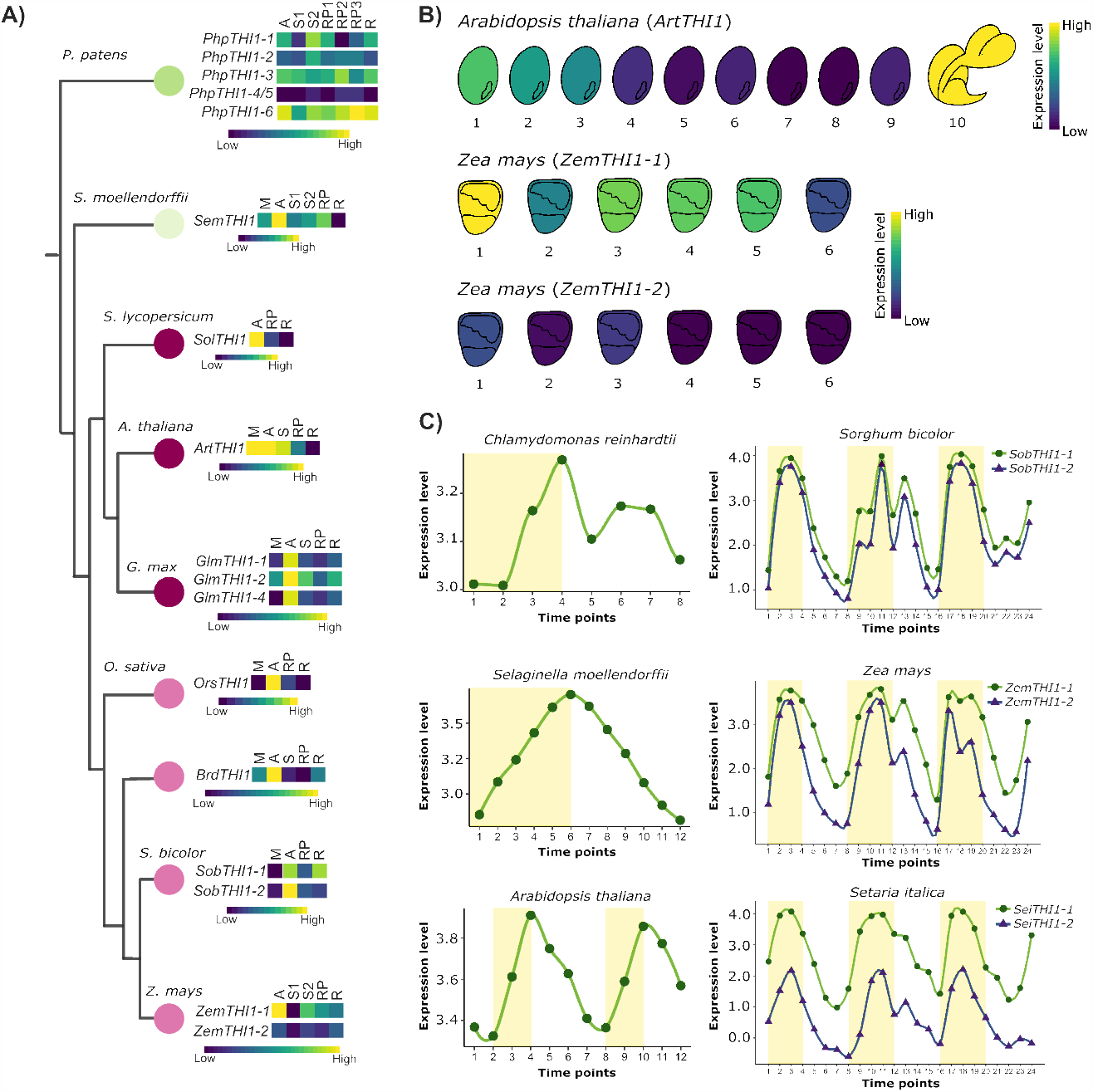
Global *THI1* expression patterns among different plant and tissues. **A)** The samples of phylogenetically related species were categorized into M-meristem; A-aerial part; S-stem; RP-reproductive tissue; and R-roots (samples detailed information are listed in Table S7). The color scale indicates the degree of expression (low to high). **B)** Schematic representation of *A. thaliana* and *Z. mays* seed development stages and *THI1* homologs expression. The colors show a normalized expression level log(TPM+1). **C)** Analysis of diurnal THI1 expression pattern. Transcript expression level was obtained from RNAseq datasets (see Table S7). The shaded areas show light phases.

Based on expression atlases, our analysis revealed distinct expression profiles of *THI1*. Out of the 95 expression signals observed for *THI1*, which includes paralogs and various tissues, it is remarkable that the “aerial part” tissue exhibited the highest expression level compared to all other samples across all observations. This finding emphasizes the significance of the “aerial part” tissue in the context of *THI1* gene expression in the study as have been reported in other studies (Ribeiro *et al*. 1996; Papini-Terzi *et al*. 2003; Ribeiro *et al*. 2005; Mangel *et al*. 2017; Joshi *et al*. 2020; Dias *et al*. 2023). The heatmap visualization provides a comprehensive overview of *THI1* expression across the samples, highlighting potential distinctions between tissues and paralogs. In addition to identifying a consensus expression pattern of *THI1* homologs where photosynthetic tissues exhibit higher expression levels than in non-photosynthetic tissues, we can notice the expression dominance of one *THI1* paralog compared to the others, such as *PhpTHI1-6* (Pp3c24_10800; *P. patens*), *GlmTHI1-2* (Glyma.20G142000; *G. max*), *SobTHI1-2* (Sobic.002G384400; *S. bicolor*), and *ZemTHI1-1* (Zm00001d044228; *Z. mays*).

Additionally, we explored the expression data from seed development stages of *A. thaliana* and *Z. mays*, guided by the GO terms highly associated with the promoter region (Asymmetrical cell division [GO:0008356], Metabolic process [GO:0008152], and Positive regulation of DNA-templated transcription [GO:0045893]) (see Figure 6C and Figure 7B). These GO terms inherently relate to the coordinated integration of genetic, metabolic, and physiological pathways during critical processes such as cell division, enlargement and accumulation of storage metabolites, which processes underlie cell differentiation and the growth of various seed compartments (Dante *et al*. 2014).

Although transcription analysis doesn’t fully answer the thiamine storage regulation, we can see an apparent accumulation of *THI1* transcripts in the seed’s initial stage, which are markable by morphogenesis, cell divisions, and differentiation of embryo (Verma *et al*. 2022) (Figure 7B). This set of cellular modifications precedes the process of storage and accumulation of metabolites and is essential for seed maturation (Su *et al*. 2021). Our findings show a prominent expression peak of *THI1* in seed during embryo formation where typical asymmetrical divisions occur, followed by a reduction over time for both species when reaching full desiccation and maturation. Intriguingly, the auxotrophic mutant *tz-201* in Arabidopsis requires thiamine supplementation for emerged plantlet growth (Papini-Terzi *et al*. 2003), and a mutant strain in *L. japonicus* exhibits immature seeds with reduced seed weight (Nagae *et al*. 2016). Collectively, these findings support the notion that thiamine biosynthesis, mediated by *THI1*, plays a pivotal role in the early stages of seed development.

After extensive research, we have successfully outlined the expression patterns of the *THI1* gene across a comprehensive sampling of organisms, encompassing various stages of the diurnal cycle. Our study involved the examination of *Chlamydomonas reinhardtii, Selaginella moellendorffii, Arabidopsis thaliana, Sorghum bicolor, Zea mays*, and *Setaria italica* (as illustrated in Figure 7C). Plant biology has had significant discourse regarding the intricate relationship between temporal factors and metabolic homeostasis. This discussion revolves around the fundamental aspects of plant health and the contributions to fitness, particularly in the plant’s capacity to dynamically regulate the balance between metabolite supply and demand (Fitzpatrick and Noordally 2021). In our investigation, we embarked on a quest to identify, across diverse plant groups, patterns associated with temporal regulation of this critical thiamine precursor gene. Our results show a clear expression pattern coordinated by the diurnal cycle in all species we analyzed (Figure 7C). Maize, Sorghum, Setaria, and Arabidopsis share the presence of “Rhythmic” and “Circadian” motifs not present in Selaginela and Chlamydomonas but all share the presence of “Response to stimulus” motif (as shown in Figure 6C). In this case, we emphasize the importance of combining different approaches in studying and characterizing genes. *THI1* is an example that the biological function of a gene throughout plant evolution can be conserved and maintained even under mutational pressure from regulatory regions.

## CONCLUDING REMARKS

Vitamin B1 is an essential co-factor to the primary metabolic pathway, and its synthesis depends on the conjugation of pyrimidine and thiazole rings. To date, only a few Archaea and Eukaryotes synthesize thiamine to supply to most remainder auxotrophic organisms. Fungi and plants are the two key eukaryotic lineages that supply most of the thiamine to animals and all other organisms. The Archaea and Eukaryotes thiazole synthesis depends on the activity of a highly conserved protein carrying a THI4-containing domain with addition of an N-terminal region in the Eukaryotic lineage. Phylogenetic analyses based on 566 curated genomes support a single origin for the *THI4*-containing gene. However, multiple specializations targeting the protein to organellar compartments and gene regulation at transcriptionally or post-transcriptionally levels are identified. Land plants reveal the presence of riboswitch regulation in some chlorophyte and streptophyte algae as well as embryophytes with the exception of Gymnosperms and Angiosperms. The genomic region containing THI1 in angiosperms is syntenic across all lineages studied whereas in the remainder lineages, the genomic location is more diverse. Two conserved motifs present at the promoter region are present in the plant lineage. In addition, 408 transcriptome data analysis, reveals conserved expression patterns in aerial parts, early seed development, and response to light/dark cycles.

## DATA AVAILABILITY

All the samples used for the RNAseq analysis are available in eFP Plant BAR and Expression with respective references (see Table S7). The Bioproject ID of RNAseq dataset are available in NCBI from *S. bicolor* (PRJNA217523), *C. reinhardtii* (PRJNA264777), *G. max* (PRJNA238493), *S. moellendorffii* (PRJNA287059), *S. italica* (PRJNA77795 and PRJNA73995) and *Z. mays* (PRJNA323555). The RNAseq dataset also collected from Gene Expression Omnibus is available under accession number from *S. moellendorffii* (GSE64665) and *O. sativa* (GSE6893 and GSE6901). The accession numbers from *P. patens* (E-MTAB-3069), *B. distachyon* (E-MTAB-5491), *A. thaliana* atlas (E-MTAB7978), *S. moellendorffii* (E-MTAB-5491) and *V. vinifera* (E-MTAB-8758 and E-MTAB-8756) Microarray data are available in the Array Express database (http://www.ebi.ac.uk/arrayexpress/). The samples’ details can be found in the Supplementary Tables S7. Codes for optimization of the analyses have been deposited in GitHub (https://github.com/leonardonaoki/THI1Utils).

## ACKNOWLEDGMENTS

The authors are grateful to Regina Miura for her motivation and curiosity about plant evolution and for encouraging us to move forward with this study because science and education transform lives and people. The authors are grateful to Dr. Tatiana Caroline Silveira Correa for technical assistance to the GaTE-Lab and total care and attention with cakes to each excellent result. We would also like to thank Luis G. O. Dantas for the careful statistical review and support with the R code. We want to thank Leonardo Naoki Higa for his help with various codes and the necessary expertise in programming that facilitated our searches. We want to thank Dora Takiya Bonadio for her kind help with choosing friendly color palettes and figure adjustments discussion.

## FUNDING

Financial support was obtained from grants FAPESP 2016/17545-8 and CNPq 308197/2010-0 to Marie-Anne Van Sluys. Henrique Moura Dias FAPESP 2019/08239-9 and FAPESP 2022/16208-9. Naiara Almeida de Toledo FAPESP 2022/09923-3 and CAPES 88887.900872/2023-00. The funders had no role in study design, data collection, analysis, publication decision, or manuscript preparation.

## AUTHOR CONTRIBUTIONS

HMD conceived and designed the bioinformatic approaches, analyzed the data, prepared figures and tables, authored or reviewed article drafts, and approved the final draft. NAT analyzed the data, prepared figures and tables, reviewed the article, and approved the final draft. RVM and JCS reviewed the supplementary analysis and article drafts and approved the final draft. MAVS conceived the project, discussed results, authored and reviewed drafts of the article, and approved the final draft.

## CONFLICTS OF INTEREST

The authors declare they have no competing interests.

## SUPPLEMENTARY DATA LIST

The following supplementary materials are available in the online version of this article.

- **Supplementary Table S1:** *THI1* homologs retrieved from Databases throughout major groups. The sequences were obtained using the sequence from *Arabdopsis thaliana THI1* (AT5G54770), the homologs were from the groups Eukarya (represented by plants and yeast), Archaea, and Protists, respectively. Each group’s phylogenetic information (Superkingdom, Kinkdom, Subkindom, Phylum, Clade, Class, and Order) is available on the NCBI taxonomy Browser. All the sheets have the information on the name of the organism, the gene ID from each database, the ID we used for our analysis, and the first letter of each nickname is representative of the letter of the group (Fungi, Archaea, Oomicota and Alveolata), underscore followed with the name of species. In plants, our focus group, we adopt other nomenclature for the genes, the two first letters of the genus followed by the first letter from the name species. We also added information about the location of the gene in the chromosome, genomic sequence length, CDS, protein size, intron and exon number, and transcript splicing sites.
- **Supplementary Table S2:** *THI1* homologs from PhycoCosm. *THI1* Homologs from groups Chlorophyta and Streptophyta supplementing the previous search after the first was absent of some representative species, these sequences enlightened our evolutionary analysis. The columns are classified as Class, Organism, Hit, Score, E-value, Alignment Length, % Hit Identity, Query Name, Query Start, Query End, Hit Name, Hit Start, Hit End, Dataset of all 68 new *THI1* homologs among 41 species of green algae (Chlorophyta) 16 sequences distributed in ten species of Streptophyta algae.
- **Supplementary Figure S3:** *THI1* syntenic blocks summary. The visualization of each organism’s syntenic genes of *THI1* is available in the Plaza Synteny plot tool. To generate the *THI1* synteny, we used the sequence of *Arabidopsis thaliana* (AT5G54770) as reference and homologs gene family ID HOM05D002858. The columns are classified in Phylum, Clade, Class/order, number of Syntenic block, Species, and the accession numbers of the genes numbered as 1 to 11, centering by *THI1* homologs. A summary manually curated, including the direction of each gene, is represented in Figure 5.
- **Supplementary Table S4:** Presence of TPP-Riboswitch in plants. Identification of the family TPP-riboswitch (THI-box element; RF00059) within the *THI1* homologs. The input used was the complete coding sequence (CDS) in the Sequence search tool of Rfam database and similar sequences from RNAcentral. The information is organized in Organism name, ID database, ID gene, ID transcript, Database we retrieved the sequence, Sequence type, Family of Riboswitch (Rfam), Accession number, Start, End, Bit score, E-value, Strand. In light red are the sequences that couldn’t match in Rfam database (Kalvari *et al*. 2021). In the case of *Klebsormidium nitens*, the genomic sequence was unavailable and the only database accessible when we conducted the analysis.
- **Supplementary Table S5:** Summary of promoter analysis. A summary of the species we selected to analyze the promoter region. Including the presence/absence of the 19 Motifs in 1,500 bp upstream to ATG from *THI1* homologs, and the transcription factors (TF) of all Motifs present in this study, more information about TFs is available in Table S6 and Figure 6.
- **Supplementary Table S6:** Motifs retrieved from MEME Suite. Information about transcription factors (TF) present in motif 1 to 19, named as MEME from MEME Suite tool classified in Link according TOMTOM tool. The p-value, Name, Matrix ID, Class, Family, Collection, Taxon, Species, Data Type, Validation, Uniprot ID were selected of the sequences with statistical support of p-value ≥0.05. The Uniprot ID were used to find the Gene Ontology (GO) terms.
- **Supplementary Table S7:** RNAseq datasets compilation. All samples of RNAseq were retrieved from eFP Plant BAR and Expression Atlas of *Chlamydomonas reinhardtii, Physcomitrium patens, Selaginella moellendorffii, Oryza sativa, Sorghum bicolor, Glycine max, Medicago truncatula, Arabidopsis thaliana, Brachypodium dystachion, Zea mays* e *Solanum lycopersicum*. The columns are divided into Species, Lineage, Database, *THI1* Homolog, Samples, Expression value, and Reference Article. We added a diurnal cycle RNAseq dataset of grasses from Lai *et al*. (2021) The visual representation of these data is in Figure 7.

## FIGURES

**Figure S1.**
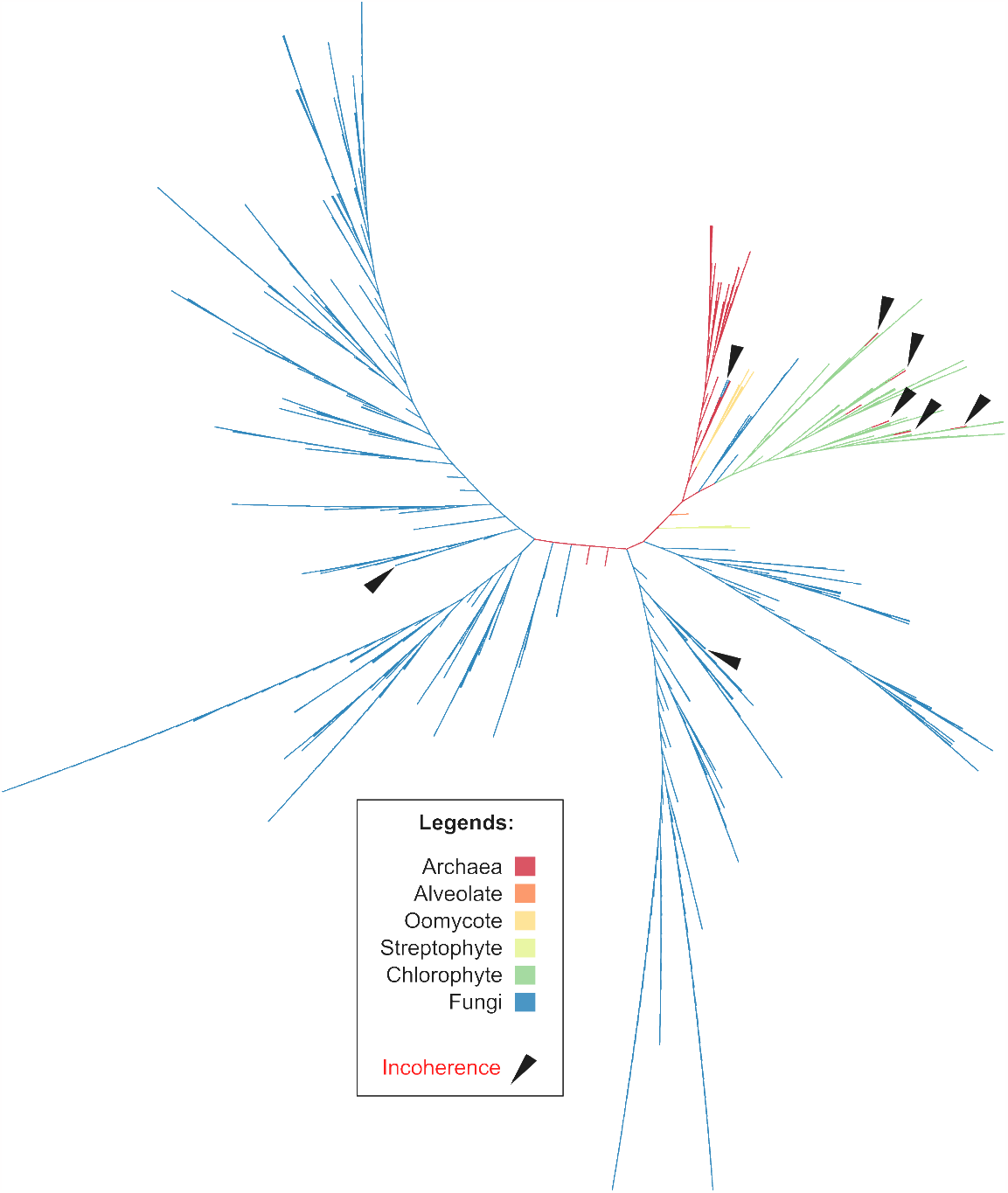
Phylogenetic tree of the THI4 protein domain (PF01946). Phylogeny of out 709 analyzed sequences from 566 genomes sampled, presenting group incongruence of Archaea, Alveolate, Oomycote, Streptophyte, Chlorophyte, and Fungi. The organisms are listed in Table S1. The high conservation degree of the THI4 domain evidences the potential barcode of the N-terminal on species diversification.

## Notes

### Competing Interest Statement

The authors have declared no competing interest.

